# Structure of the portal complex from *Staphylococcus aureus* pathogenicity island 1 transducing particles in situ and in solution

**DOI:** 10.1101/2023.09.18.557803

**Authors:** Amarshi Mukherjee, James L. Kizziah, N’Toia C. Hawkins, Mohamed O. Nasef, Laura K. Parker, Terje Dokland

## Abstract

*Staphylococcus aureus* is an important human pathogen, and the prevalence of antibiotic resistance is a major public health concern. The evolution of pathogenicity and resistance in *S. aureus* often involves acquisition of mobile genetic elements (MGEs). Bacteriophages play an especially important role, since transduction represents the main mechanism for horizontal gene transfer. *S. aureus* pathogenicity islands (SaPIs), including SaPI1, are MGEs that carry genes encoding virulence factors, and are mobilized at high frequency through interactions with specific “helper” bacteriophages, such as 80α, leading to packaging of the SaPI genomes into virions made from structural proteins supplied by the helper. Among these structural proteins is the portal protein, which forms a ring-like portal at a fivefold vertex of the capsid, through which the DNA is packaged during virion assembly and ejected upon infection of the host. We have used high-resolution cryo-electron microscopy to determine structures of the *S. aureus* bacteriophage 80α portal in solution and in situ in the empty and full SaPI1 virions, and show how the portal interacts with the capsid. These structures provide a basis for understanding portal and capsid assembly and the conformational changes that occur upon DNA packaging and ejection.

## INTRODUCTION

*Staphylococcus aureus* is an opportunistic human pathogen and one of the leading causes of deaths associated with antibiotic resistance [1]. *S. aureus* encodes a large number of virulence factors, including adhesins, toxins and other proteins that enable the bacteria to bypass the host’s immune response and persist in specific environments [2]. Most of these virulence factors are encoded on mobile genetic elements (MGEs), such as plasmids, bacteriophages (phages) and chromosomal islands that are subject to horizontal gene transfer (HGT) between cells [3,4].

Tailed bacteriophages (class *Caudoviricetes*—formerly order *Caudovirales*) are the main mediators of HGT in *S. aureus*, since these bacteria are not generally transformable, and conjugation is rare [5]. These phages assemble empty precursor capsids (procapsids) from a capsid protein (CP), typically with the assistance of a scaffolding protein (SP) that acts as a chaperone for the assembly process [6,7]. These phages also encode a portal protein (PP) that forms a ring-like portal at one capsid vertex. The portal is assumed to act as the nucleus for capsid assembly, and serves as an entry and exit point for the DNA [8–10]. The portal, together with the terminase, consisting of large (TerL) and small (TerS) subunits, comprise the DNA packaging machinery of the phage [11]. Capsid maturation involves shell expansion and removal of the SP, which is followed by attachment of tails to the portal complex.

Phage 80α is a typical temperate staphylococcal siphovirus traditionally placed in the *Siphoviridae* family; currently belonging to the *Azeredovirinae* [12,13]. The 80α virion has a 63 nm icosahedral head with *T*=7 architecture, a 190 nm long flexuous tail, a complex baseplate [14–16], and a 43,864 base pair double-stranded (ds)DNA genome [17](NCBI RefSeq NC_009526.1). Phages closely related to 80α are found in a wide variety of staphylococcal strains, including MRSA strain USA300 LAC, commonly involved in community-acquired infections [18].

*S. aureus* pathogenicity islands (SaPIs) are a type of phage-inducible chromosomal islands (PICIs), MGEs that become mobilized at high frequency by so-called “helper” phages [19–21]. SaPIs are common in most strains of *S. aureus* and encode a range of virulence factors, including superantigen toxins (SAgs), adhesins and anti-coagulation factors. SaPIs are normally stably integrated into the host genome, but become derepressed and excise upon infection with a compatible helper phage (or induction of a prophage), followed by replication and packaging into transducing particles made up of phage-encoded structural proteins. 80α can act as helper phage for a number of SaPIs, including SaPI1, SaPI2, SaPIbov1 and SaPIbov5, each of which is derepressed by a different 80α-encoded protein [20].

In many cases, SaPIs redirect the assembly pathway of its helper phage to form capsids of a smaller size than that normally made by the phage [21,22]. Thus, when 80α acts as a helper for SaPI1, assembly is redirected to form capsids with *T*=4 architecture rather than the normal *T*=7 80α procapsids [23,24]. The smaller capsids are compatible with the size of the SaPI genome (typically ≈15 kbp), but are too small to package complete phage genomes (≈45 kbp), leading to strong suppression of phage multiplication. Many SaPIs encode distinct TerS proteins that specifically recognize their respective genomes [25,26]. However, the mechanism of DNA selection and the interaction between the terminases and the portal are not clear.

Capsid size redirection by SaPI1 involves two SaPI1-encoded proteins, CpmA and CpmB [24,27,28]. We previously determined the structure of 80α and SaPI1 procapsids to 3.8 Å and 3.7 Å resolution, respectively [23,24], and showed that CpmB forms an alternative internal scaffold in the SaPI1 procapsids. The role of CpmA is still not well understood. We also determined structures of mature 80α and SaPI1 capsids; however, these reconstructions only reached modest resolution (5.2 Å for 80α and 8.4 Å for SaPI1) [15]. Moreover, all previous 80α and SaPI1 reconstructions were done with icosahedral symmetry applied, and thus did not show the portal.

Here, we have used cryo-electron microscopy (cryo-EM) to determine a 2.4 Å resolution structure of the 80α PP expressed in *Escherichia coli*, where it forms a tridecameric ring. We also determined asymmetric structures of both full and empty mature capsids from SaPI1 virions. Icosahedral reconstruction yielded a map of the full capsids at 3.1 Å resolution, which was used to model the CP. Focused reconstruction was then used to resolve the portal at 3.2 Å resolution in situ, where it forms a dodecamer. Using focused reconstruction, we show details of the interaction between the portal and the capsid. These are the first structures of a portal complex from a staphylococcal siphovirus and from a SaPI transducing particle. Together, these structures provide insights into the role of the 80α PP in capsid assembly, DNA packaging, and the process of phage-induced genetic mobilization of SaPIs.

## RESULTS

### 1. Reconstruction of the 80α portal in solution

The gene (ORF42) encoding the 80α PP (gp42) was cloned into a pET21a-based expression vector with a C-terminal hexahistidine tag and overexpressed in *E. coli*, followed by purification by Ni-NTA affinity. The protein was further purified by sucrose gradient centrifugation and subjected to cryo-EM imaging, which showed the expected cartwheel-like rings with a diameter of around 160 Å and a hole in the middle (Fig. 1A). 2D classification in RELION-3 [29,30] showed that the predominant form was a 13-fold symmetric ring (Fig. 1A). No other symmetrical oligomers were observed. A total of 96,308 particles were used for the final reconstruction (Fig. 2A, B), which reached a resolution of 2.4 Å with the application of C13 symmetry, according to the gold standard Fourier shell correlation (FSC) measure (FSC=0.143) (Table S1; Fig. S1). An atomic model was generated using AlphaFold [31] as a starting point, and refined with Coot [32] and ISOLDE [33] to a model-to-map resolution of 2.5 Å (FSC=0.5) (Table S1). There was no density for the C-terminal 35 residues, which were thus omitted from the model.

**Figure 1.**
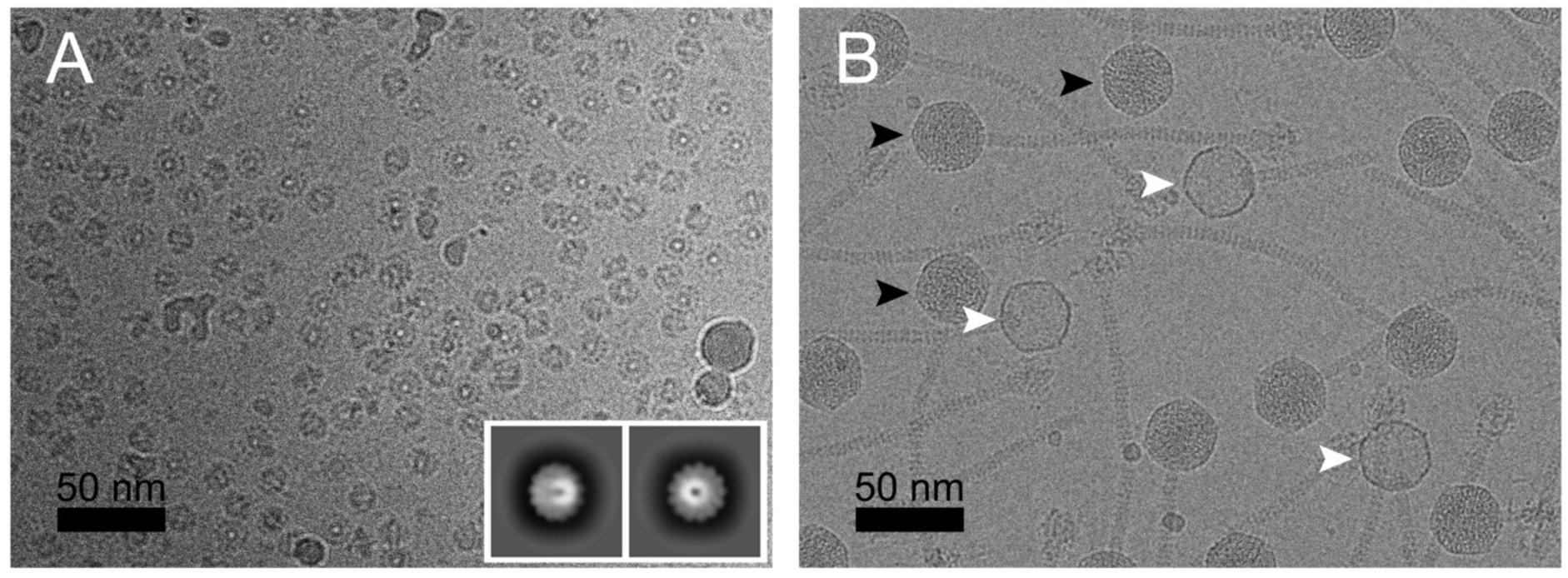
**(A)** Cryo-electron micrograph of 80α portals expressed in *E. coli*. The inset shows representative 2D classes. **(B)** Cryo-EM of SaPI1 virions produced by induction of *S. aureus* strain ST65 (80αϕι*44* + SaPI1*tst::tetM*). Examples of full and empty capsids are indicated by black and white arrowheads, respectively. Scale bars = 50 nm.

**Figure 2.**
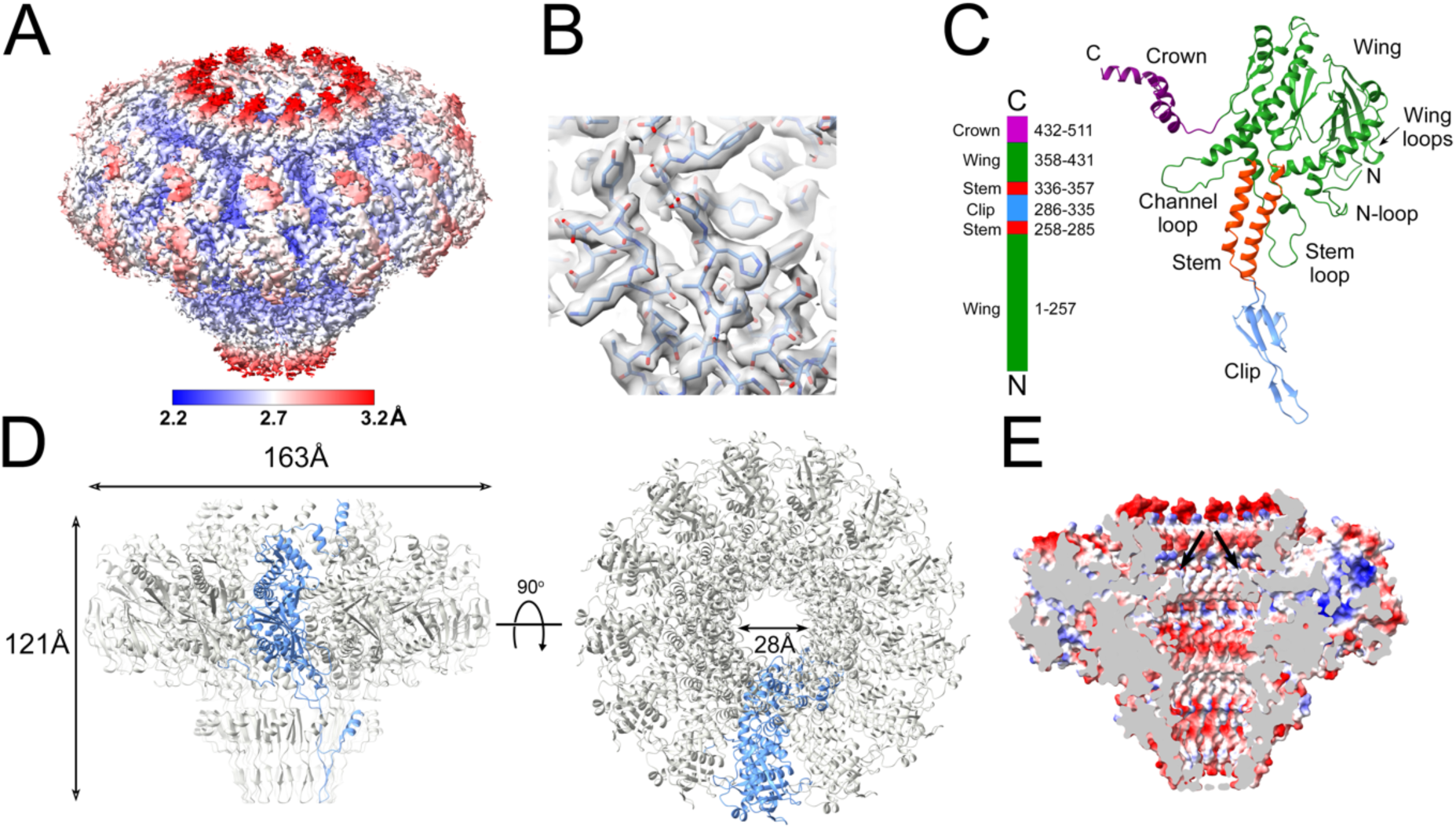
Reconstruction of portals expressed in *E. coli*. **(A)** Isosurface representation of density map, colored by local resolution according to the color bar. **(B)** Detail of density with the atomic model (residues 67–76 and 90–98) fitted in. **(C)** Ribbon representation of the PP monomer, colored by domain: wing, green; stem, red; clip, blue; crown, purple. The N- and C-termini and other structural features are indicated. A linear representation of the PP sequence is shown on the left. **(D)** Ribbon representation of the PP tridecamer, viewed from the side (left) and top (right). One subunit is shown in blue, the other subunits are gray. **(E)** Electrostatic potential cutaway surface of the solution portal, colored by charge from red (negative) to blue (positive). The channel loop constriction is indicated (arrows).

The PP monomer resembles those of other bacteriophages [8,9] and contains four (sub-)domains: *wing* (residues 1–257 and 358–431), *stem* (258–285, 336–357), *clip* (or *stalk*; 286–335) and *crown* (432–480) (Fig. 2C). Thirteen copies of PP form a 163 Å wide, 121 Å tall 13-fold symmetrical ring with a 28 Å wide central channel (Fig. 2D).

The wing domain forms the bulk of the PP and consists of an intertwined α/β fold (Fig. 2C). Residues 357–373 of the wing domain constitute the “channel loop” that was shown in other systems to interact with the DNA inside the central channel [9] (Fig. 2C). The channel loop forms the narrowest point in the channel with a crosswise distance between opposite Gly 369 residues of 28.5 Å (Fig. 2D). The N-terminus was ordered from residue 1 of the PP sequence, and forms a loop (the “N-loop”) on the outside of the wing domain (Fig. 2C). The β-sheet in the wing domain forms two loops on the outside of the portal that we refer to as the “wing loops” (Fig. 2C). Another loop (the “stem loop”) at the bottom of the wing domain protrudes outside the stem domain.

The stem domain consists of two 30 Å long α-helices (Fig. 2C). The second α-helix (residues 334- 355) lines the central channel like a cone with a closest approach of 34.0 Å between residues Gln 340 at the bottom of the helix. The inner lining of the channel is predominantly negatively charged presumably presenting a “slippery” surface for the DNA (Fig. 2E).

The clip domain consists of a 44 Å long β-hairpin with a small α-helical loop insertion (Fig. 2C). The hairpin interacts with the head-to-tail connector (see below) and most likely connects with the terminase during packaging, as described for P22 [34], T4 [35] and HK97 [36].

The C-terminal crown domain consists of three α-helices that form a dome-like structure at the top of the portal where it can interact with the DNA inside the capsid (Fig. 2C; see below). Part of the missing C-terminal 35 residues is predicted by AlphaFold [31] to form an α-helical extension to the crown domain α-helix.

### 2. Reconstruction of the SaPI1 virion

SaPI1 virions were produced by induction of the 80α lysogen ST65, which harbors a SaPI1 *tst::tetM* element [23]. (This strain also has a deletion of 80α ORF44, which we previously showed to be detrimental to phage viability, but had no effect on particle assembly or SaPI1 transduction efficiency [37].) The SaPI1 virions were purified on CsCl and sucrose gradients and subjected to cryo-EM imaging (Fig. 1B). The majority of the capsids appeared to be filled with DNA, but a smaller subset (≈20%) were empty (Fig. 1B). After 2D classification, 55,513 full and 13,774 empty particles were selected for separate 3D refinement without the application of symmetry (C1).

Both reconstructions displayed the expected *T*=4 capsid shell and a unique vertex with the tail attached (Fig. 3A-C). The portal was clearly visible inside the empty capsid (Fig. 3B), while in the full capsid, the portal was obscured by layers of DNA (Fig. 3C). Postprocessing of the maps yielded resolutions of 4.7 Å and 5.4 Å for full and empty capsids, respectively (Table S1; Fig. S1), but while the CP was clearly resolved, the portal density remained uninterpretable, presumably because the symmetry of the capsid dominated the alignment of the particles. Three-dimensional classification was unable to resolve this ambiguity, and the portal structure was instead resolved by focused reconstruction (see below).

**Figure 3.**
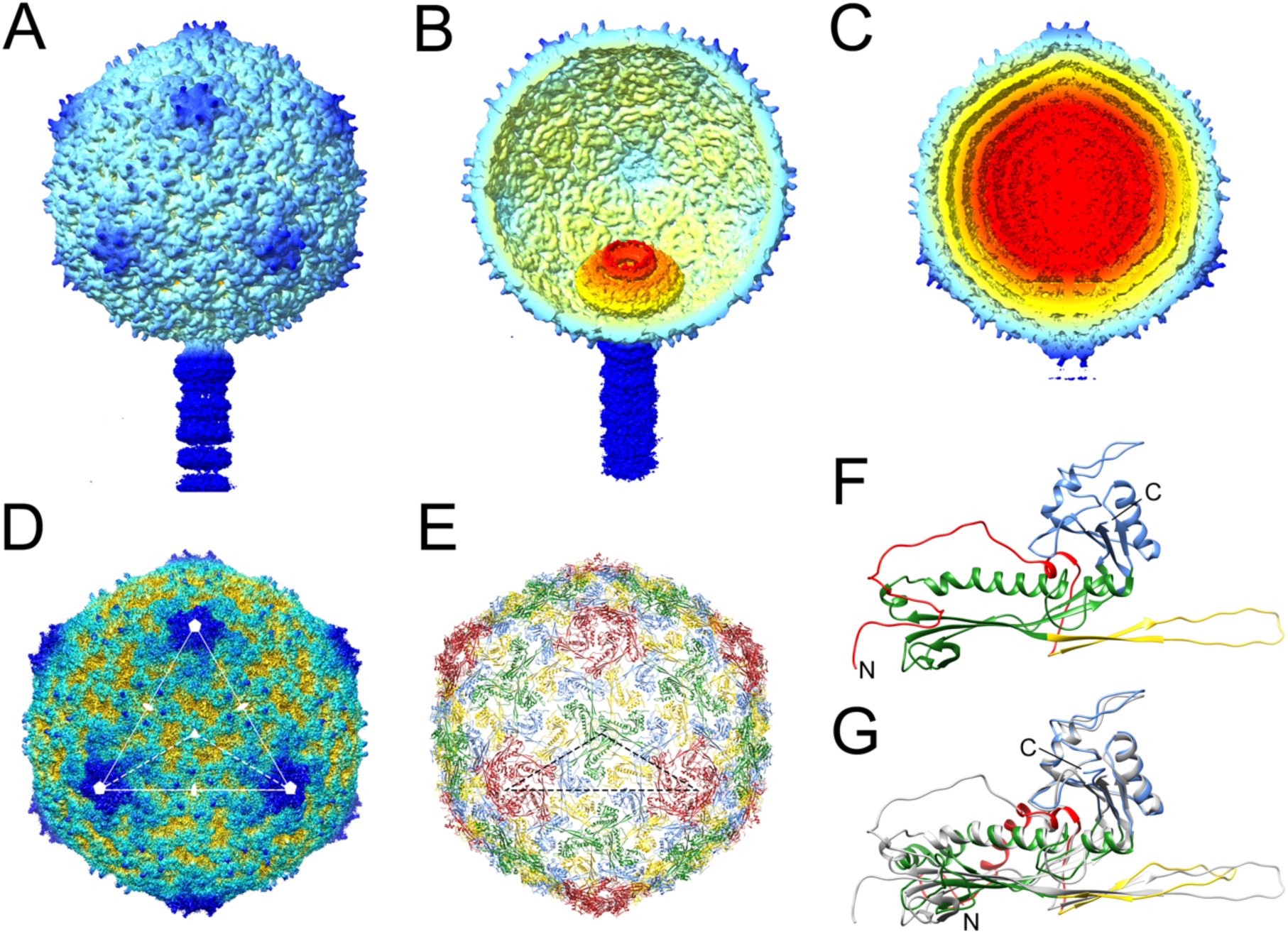
Reconstruction and modeling of SaPI1 virions. **(A)** Isosurface representation of the asymmetric reconstruction of the empty SaPI1 capsid. **(B)** Cutaway view of the empty capsid reconstruction, tilted to show the portal inside. **(C)** Cutaway view of the asymmetric reconstruction of the full capsid, viewed as in (A). **(D)** Icosahedral reconstruction of the full capsid, viewed down an icosahedral threefold axis. One triangular face with symmetry axes is indicated. The dashed triangle shows an icosahedral asymmetric unit. **(E)** Atomic model of the CP in the icosahedral capsid, oriented as in (D) and colored by subunit: A, red; B, blue; C, green; D, yellow. The dashed triangle shows the icosahedral asymmetric unit as in (D). **(F)** Ribbon representation of CP monomer (subunit B), colored by domain: N-arm, red; E-loop; yellow; P domain, green; A domain, blue. N- and C-termini are indicated. **(G)** Superposition of CP subunit B from the virion (gray), superimposed on the same subunit from the SaPI1 procapsid (PDB ID: 6B23) [24], colored by domain as in (F).

We carried out 3D refinement of the full and empty capsids with the application of icosahedral symmetry, which reached 3.1 Å and 3.2 Å resolution, respectively (Fig. 3D; Table S1; Fig. S1). The portal complex is not visible in the icosahedral reconstructions due to averaging. However, we used this higher-resolution reconstruction (full capsid) to model the CP. The previous structure of the SaPI1 virion only reached a resolution of 8.4 Å and did not allow atomic modeling of the CP [15]. In the 3.1 Å resolution structure, CP residues 16–313 could be modeled with confidence (Fig. 3E,F). (The first 14 residues are not present in the protein due to cleavage by the Prp protease [27,38].)

The icosahedral SaPI1 capsid has a diameter of ≈500 Å and consists of 240 copies of CP (gp47) arranged with *T*=4 symmetry (Fig. 3E). (In the asymmetric structure, five subunits at one fivefold vertex would be replaced by the portal.) There are thus four copies of CP in the asymmetric unit, designated A–D (Fig. 3E). The CP has the expected HK97-like capsid fold described previously, consisting of an N-arm (residues 15–67), an E-loop (68–104), a P-domain (105–167 and 248– 307), and an A-domain (168–247 and 308–324) (Fig 3F).

Around the icosahedral fivefold and twofold axes, the A-domains interact to form pentameric (five A subunits) and hexameric (two copies each of B, C and D subunits) rings, respectively (Fig. S2A,B). In the A subunit, a six amino acid loop (residues 211–216) around the fivefold axis was disordered, apparently due to steric hindrance induced during capsid expansion (Fig. S2A). Other intra-capsomeric contacts are made between the E-loops and the adjacent P-domains, as previously described (Fig. S2C,D). Inter-capsomeric contacts are formed primarily by the P-domains, which include a P-loop (residues 268–284) that forms trimeric “turrets” at the icosahedral threefold (made from three C subunits) and quasi-threefold (made from A, B and D subunits) axes [15,24] (Fig. S2C,D). The N-arms of the same subunits and the tips of the E-loops from adjacent subunits wrap around these turrets, presumably reinforcing the threefold interactions between capsomers (Fig. S2C,D).

Superposition of the CP subunit from the SaPI1 virion to that in the SaPI1 procapsid (PDB ID: 6B23) [24], revealed the conformational differences that occur during capsid maturation. These differences include a flattening of the E-loop and a straightening of the P-domain helix α3 (the “spine” helix), similar to what we previously described for 80α [14,15] (Fig. 3G). Moreover, in the SaPI1 procapsid, the N-arms point towards the inside of the capsid where they interact with CpmB and SP via the N-arm α-helix (α1) [24], whereas in the virion, the N-arms are extruded to the outside of the capsid and lack the α-helix (Fig. 3F,G). This rearrangement is partly accommodated by a cis-proline at residue Pro 26, which induces a sharp turn in the N-arm. In the procapsid, the N-arm could only be modeled from Pro 26, while the virion N-arm could be modeled from residue 16, just after the Prp cleavage site, suggesting that the N-arm in the virion is stabilized by interactions with the turrets on the outside of the capsid (Fig. S2C,D).

### 3. Structure of the portal in situ

As mentioned above, the portal could not be resolved in the asymmetric reconstructions of either the full or empty capsids. In order to produce a high-resolution structure of the portal in situ, the portal complex, including part of the tail, from the full and empty capsids was subjected to focused reconstruction after signal subtraction to remove the contribution of the capsid. As the portal clearly displayed twelvefold symmetry, refinement in C12 was carried out, reaching a resolution of 3.2 Å and 3.5 Å for full and empty capsids, respectively (Fig. 4A; Table S1; Fig. S1). Atomic models for the dodecamer were generated for both. In addition to missing density for the C-terminal 29 residues—slightly less than in solution—density for the N-terminal 15 residues was missing from both in situ PP models, apparently resulting from interactions with the fivefold symmetric capsid (see below).

**Figure 4.**
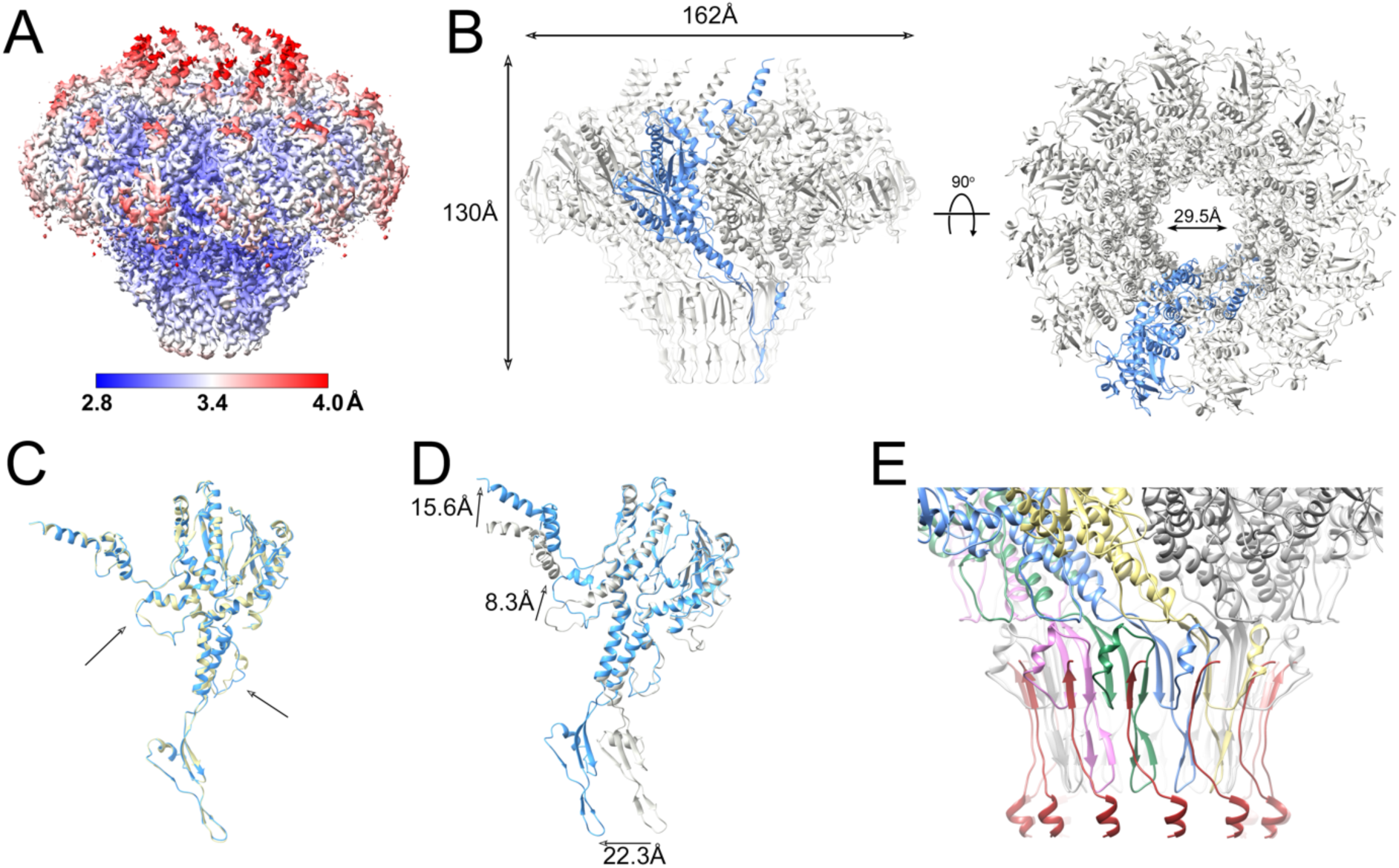
Focused reconstruction of the portal complex in situ. **(A)** Isosurface representation of the C12 focused reconstruction from the full SaPI1 capsid, colored by local resolution, according to the color bar. **(B)** Atomic model of the portal dodecamer, viewed from the side (left) and top (right). One subunit is shown in blue; other subunits are gray. **(C)** Atomic model of PP in situ from the full capsid (blue) superimposed on the empty capsid (tan). Minor differences in the channel loop and the stem loop are indicated (arrows). **(D)** Superposition of the PP monomer in situ (blue), superimposed on the PP monomer in solution (gray). The movements in the clip domain, channel loop and crown domain are indicated by the arrows. **(E)** Interaction between the head-to-tail connector C-termini (dark red) with the PP. Four PP subunits are colored in yellow, blue, green and pink to highlight the β-sheets comprised of strands from two PP subunits and the connector.

The *in situ* portal dodecamer has a similar outer diameter of 162 Å to the solution portal tridecamer, but is taller (130 Å), and has a slightly wider central channel (29.5 Å) (Fig. 4B). Few differences were observed between the PP in the full and empty capsids, except for minor alterations in areas where the density was poor, such as the stem loop, or in the channel loop where the PP interacts with the DNA (Fig. 4C). In particular, we did not observe any significant differences in the crown domain that could reflect an interaction with the genome, other than the four additional ordered residues.

There are several differences in the PP conformation between the solution tridecamer and the in situ dodecamer: Apart from the missing N-terminal residues in the in situ portal, there is a 22.3 Å shift of the clip domain and a 15.6 Å shift of the crown domain α-helices in the in situ dodecamer (Fig. 4D). In addition, there is a rearrangement of the channel loop, mediated by a kink around Gly 382 in a long α-helix in the wing domain, leading to an 8.3 Å shift in the loop itself (Fig. 4D).

In the focused reconstruction of the in situ portal protein, there was additional density below the portal. This density corresponds to the head-to-tail connector protein, or “connector,” which forms the attachment point for the phage tail. The connector is identified as gp49 (ORF49) in the RefSeq entry for 80α (NC_009526.1). The C-terminus of the connector adds a β-strand to the clip domain of PP, forming a three-stranded β-sheet together with one strand from each of two adjacent PP subunits (Fig. 4E). The remaining density was of insufficient quality to build a complete model for the connector protein. The connector and the other structures associated with the tail will be the subject of a separate study.

### 4. Comparison with other portal proteins

The in situ structure of the 80α PP is overall similar to that of SPP1 (PDB ID: 7Z4W) [39], for which it superimposes with an RMSD = 1.2 Å for 224 out of 422 residue pairs (7.2 Å between all pairs; Table S2). The main difference between the two proteins is a longer clip domain in 80α PP and an extended β-hairpin in the stem loop of SPP1 PP (Fig. 5A). In addition, the SPP1 sequence is ≈10 residues shorter at the N-terminus, and the first 16 residues are disordered. Like 80α, the SPP1 PP forms a tridecamer in solution [40]. The solution structures of the portal proteins from SPP1 (PDB ID: 2JES) and 80α superimpose with RMSD = 1.2 Å for 170/354 residue pairs and 8.2 Å overall (Table S2). The SPP1 PP exhibits a similar shift in the clip domain between the solution and in situ structures that we observed in 80α PP, but the shift is not as pronounced as in 80α (Fig. 4D, 5B), and there is no rearrangement of the channel loop. (In the crystal structure of the SPP1 PP in solution, the β-sheet in the wing domain is missing.)

**Figure 5.**
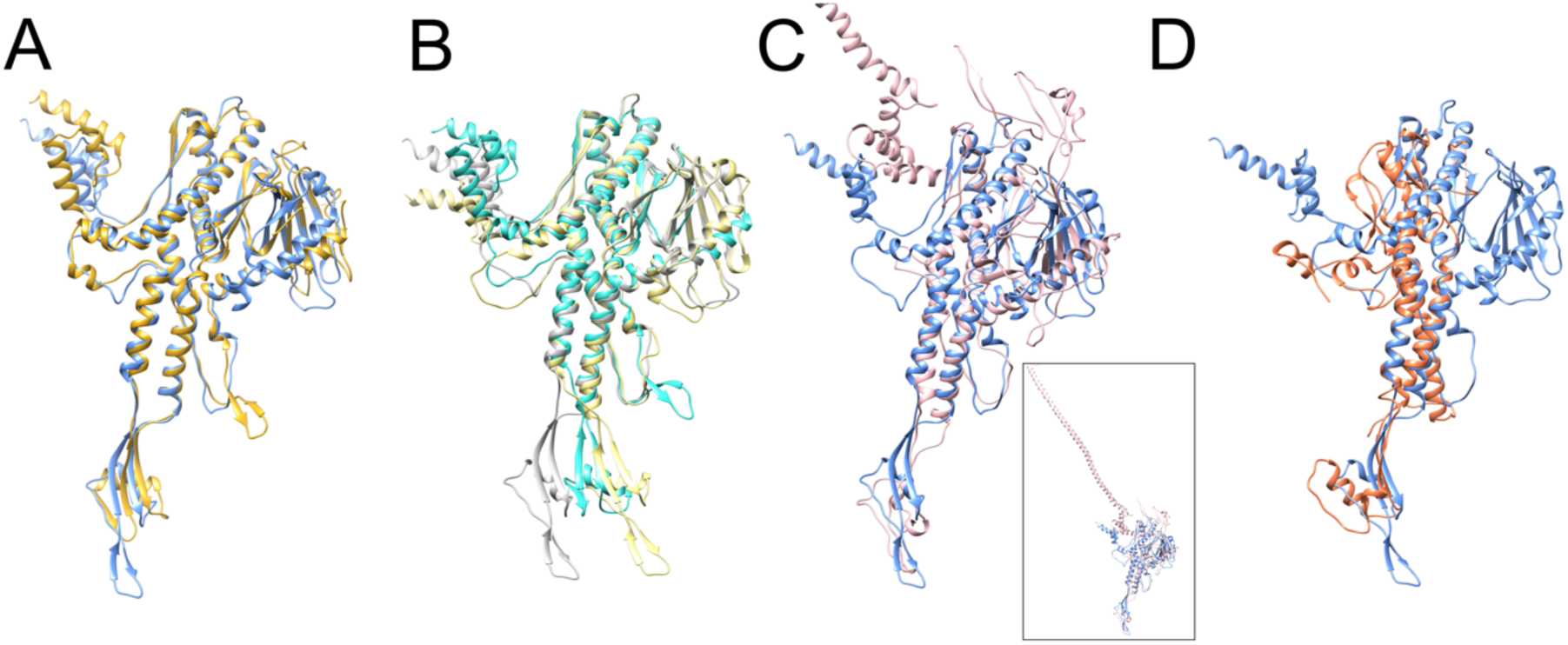
Comparison of the 80α PP with other phages. **(A)** Superposition of the 80α in situ PP (blue) with the PP of the SPP1 portal in situ (gold; PDB ID: 7Z4W) [39]. **(B)** Superposition of the 80α PP solution structure (tan) with the solution structure of the SPP1 PP (turquoise; PDB ID: 2JES)[40]. The 80α in situ PP is shown in gray. **(C)** Superposition of the 80α in situ PP (blue) with the P22 PP (pink; PDB ID: 3LJ5) [41]. For clarity, the extended barrel domain has been truncated. The entire structure is shown in the inset. **(D)** Superposition of the 80a in situ PP (blue) with the PP from phage Andhra (orange; PDB ID: 8EGR) [42].

The P22 portal protein (PDB ID: 3LJ5) is more different, with an overall RMSD of 16.1 Å compared to 80α PP (Table S2). The P22 PP sequence lacks about 40 residues at the N-terminus compared to 80α. Differences between these two proteins include two extended loops in the wing domain of P22 PP; The most striking difference, however, is in the crown domain, where the P22 PP has a C-terminal 132-residue α-helical extension that forms a 187 Å long barrel in the mature virion structure [34,41] (Fig. 5C).

Andhra is a staphylococcal phage related to ϕ29 [42]. The Andhra PP (PDB ID: 8EGR) has a much simpler structure that those of 80α, SPP1 or P22, and lacks both the β-sheet and the elaborate loops in the wing domain, and lacks the crown domain entirely (Fig. 5D), similar to ϕ29 (PDB ID: 1JNB) [43]. The overall RMSD between 80α PP in situ and Andhra is 26.0 Å (Table S2).

Unusually, the 80α PP sequence has no tryptophan residues and the portal thus lacks the tryptophan “belt” that was proposed to be important for capsid interactions in other portals, including SPP1, P22, and ϕ29 [9,44].

### 5. Interactions between portal and capsid

The C12 portal interacts with the capsid at a fivefold vertex, leading to a symmetry mismatch between PP and CP. To examine the specific interactions between the CP and PP at the mismatch interface, we used a combination of symmetry expansion and 3D classification in cryoSPARC [45] to generate an asymmetric (C1) reconstruction that included the portal assembly and part of the capsid. This reconstruction reached a resolution of 3.4 Å (Table S1; Fig. S1), allowing modeling of the PP-CP interface (Fig. 6A).

**Figure 6.**
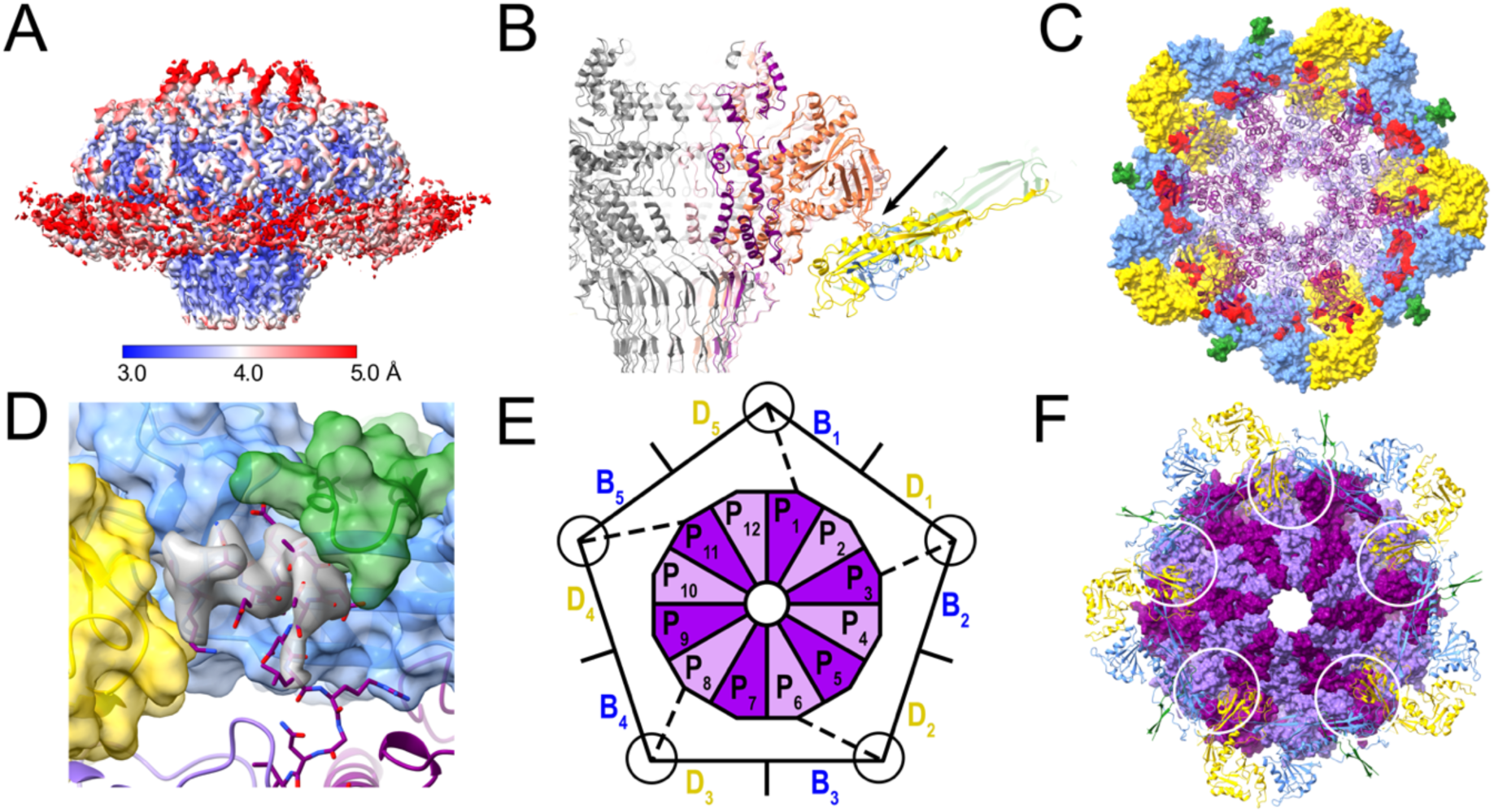
Interaction between PP and CP. **(A)** Isosurface representation of the asymmetric reconstruction of the portal-capsid interface, colored by local resolution, according to the color bar. **(B)** Slice through the ribbon representation of the atomic model of the dodecameric portal viewed from the side. Interacting CP subunits are colored by subunit, as in Fig. 3 (B, blue; C, green; D, yellow). Three PP subunits are colored pink, orange and purple. The main point of interaction with the wing loops is indicated by the arrow. **(C)** The portal dodecamer in ribbon representation, viewed from the top. Surrounding CP subunits are shown as molecular surfaces (colored as in B) with CP-SP contacts highlighted in red. **(D)** Detail of the trivalent interaction between the PP N-terminus (stick representation with purple backbone) and neighboring CP subunits, colored as in (B). The CP subunits are shown as molecular surfaces, colored according to CP subunit. Extra density from the C5 symmetrized asymmetric reconstruction, attributed to PP, is shown in gray. **(E)** Schematic diagram showing the asymmetric interaction between the PP dodecamer (P_1_–P_12_) and the ten surrounding CP subunits (B_1_–B_5_ and D_1_–D_5_), viewed from the bottom. The N-termini of five PP subunits that interact with two CP subunits are indicated by the dashed line, and the circle represents the trivalent interaction. **(F)** Atomic model of the portal dodecamer represented as a molecular surface (alternating subunits colored dark and light purple) and viewed from the bottom, with surrounding CP subunits in ribbon representation. The trimeric interactions forming the asymmetric turrets are circled.

Due to the symmetry mismatch, interactions between PP and CP vary with subunit. The PP wing loops (residues 193–203 and 222–230) interact with the P-domains of CP subunits B and D (Fig. 6B), with a surface area of interaction varying from 290–642 Å^2^ (Fig. 6C).

In the icosahedral capsid reconstruction, the turrets at the quasi-threefold axes are comprised of P-loops from CP subunits A, B and D (Fig. S2D). Since the portal replaces a CP pentamer, there are no A subunits to complete the turrets surrounding the portal, which are made only from B and D subunits (Fig. 6D). However, after modeling of the B and D subunits, there was additional density in the turrets that could be attributed to the N-termini of PP (Fig. 6D). Thus, the N-termini from PP subunits 1, 3, 6, 8 and 11 interact with the P-loops from CP subunits B and D to form pseudo-trimeric turrets that replace the turrets made from three P-loops in the rest of the capsid (Fig. 6E,F). The N-termini from the remaining seven PP subunits as well as the residues spanning the gap between the turrets and the portal are apparently not ordered and were not seen in the map. This asymmetric organization explains the absence of the N-termini from the C12 reconstruction of the in situ portal (Fig. 4).

## DISCUSSION

The portal is the central platform upon which a dynamic complement of other structural proteins is assembled at various stages in the phage life cycle. Several structures of portal proteins in solution or in situ have been determined to date, either by X-ray crystallography or by cryo-EM (Reviewed in [8–10]). From staphylococcal phages, the only prior portal structures were those of P68 [46] and Andhra [42], which belong to a distinct lineage of podoviruses (family *Rountreeviridae*) with a unique replication and packaging strategy akin to that of *Bacillus* phage ϕ29. The current structure thus represents the first PP structure from a staphylococcal siphovirus. Portal protein structures from different phages are generally conserved, reflecting their evolutionary relationship. The α-helical stem domain, which forms the main channel of the portal, is the most highly conserved. The wing subdomains tend to be more variable, and there is considerable variation in the crown and clip domains (Fig. 5).

Portals in situ are invariably dodecamers [8,9]. However, portals produced after overexpression in *E. coli* frequently adopt other oligomeric states. Tridecamers, like those we observed here, were also found for *Bacillus* phage SPP1 [40,47,48]; *Salmonella* phage P22 formed undecamers and dodecamers [48], while a distribution from 11- to 14-mer was observed for herpes simplex virus 1 [49]. Asymmetric oligomers are also observed, including spiraling forms or rings appearing to be missing one or more subunits [50,51].

Why do portals form these “unproductive” oligomers, if only dodecamers are useful for capsid assembly? It seems unlikely that tridecamers are formed initially and that one subunit is subsequently removed before or during incorporation into nascent capsids.

It may be that portal proteins exist in a dynamic equilibrium between monomers and oligomers, and that tridecamers are the most stable oligomer at the high intracellular concentrations that exist during overexpression. During capsid assembly, incorporation of dodecamers into procapsids could keep the PP concentration below the level where the tridecamers form and thereby drive the equilibrium towards dodecamers. The oligomeric state of the portal could also be regulated through interactions with other structural proteins, including SP and CP. In P22, PP oligomerization *in vivo* was shown to be driven by SP [52,53], although it is not clear whether this interaction affected the oligomeric state per se.

Capsids can be produced in vitro and in vivo without portals; however, during phage production, the portal is assumed to constitute the nucleus for capsid formation. This ensures that capsids contain one and only one tail, and is critical to the integrity of the DNA packaging process [9]. Our observation that the PP N-termini form pseudo-trivalent turrets together with P-loops from two CP subunits suggests that this might be the first step in nucleus formation, since trimerization is the key step that brings hexamers and pentamers together to form a complete shell. Most likely this nucleus also includes SP: preformed portal oligomers in complex with SP constituted the nuclei for assembly in P22, whereas PP monomers were not incorporated [52]. An interaction between SP and PP was also demonstrated for ϕ29 [54] and for the scaffolding δ domain of CP in HK97 [55]. While it has not yet been demonstrated experimentally, it is likely that the 80α PP also interacts with the SP during capsid assembly.

The assembly pathway of 80α is redirected by SaPI1 to form the *T*=4 capsids that we observe here. This redirection is dependent on the SaPI1-encoded proteins CpmA and CpmB [27,28]. CpmB forms an alternate, internal scaffold in SaPI1 procapsids [23,24]; however, the role of CpmA remains mysterious. One possibility is that CpmA interacts with PP to establish an alternative assembly nucleus. This PP-CpmA nucleus could cause a change in the angle of the nascent shell upon addition of CP subunits, either directly or through interactions with CpmB.

During DNA packaging, the terminase complex docks with the portal. The large terminase (TerL), or packaging ATPase, exists as a pentamer [36,56,57]. The mismatch between the fivefold symmetric terminase and the dodecameric portal may be important for terminase function, since TerL needs to undergo dynamic changes during packaging, and to ensure that TerL can be removed once packaging has been completed and/or the DNA has been cleaved. This interaction is mediated by the clip region, as was shown structurally for HK97 [36] and mutationally in T4 [35]. The shift in the clip domain observed between structures of PP in solution and in situ observed here (Fig. 4D) and in other systems, such as SPP1 [39,40](Fig. 5A,B), could reflect mobility required to accommodate this dynamic interaction.

Phages that package headfuls of DNA require a mechanism to detect the fill level of the capsid, in order to signal to the terminase to cleave the DNA and detach from the capsid. The crown domain extension that forms a long α-helical barrel was proposed to act as a pressure sensor for the DNA fill level in P22 [34]. The portal is presumed to transmit this information to TerL via conformational changes. Consistent with this, most of this barrel was disordered in the P22 procapsid, where the portal itself was also asymmetric. Conversely, the lack of a crown domain in the portal proteins from picoviruses like ϕ29 and Andhra presumably reflects the fact that these phages package exact genomic units of DNA and thus do not need a fill level sensor. Nevertheless, many phages that do package their DNA via a headful mechanism, including 80α and SPP1, do not have the long α-helical barrel, and we observed no major differences in the crown domain between the solution and in situ portals or between the empty and full capsids that could reflect an involvement in DNA packaging (Fig. 4C). Apparently, these phages are able to sense and transmit the capsid fill level information by a different mechanism. It should be noted, however, that we do not have the structure of the 80α portal in situ in the procapsid, where it might have a different structure from the virion.

Upon completion of packaging, the terminase is released, and the dodecameric connector (sometimes called “adaptor” [39,58]) is attached to the portal to allow for subsequent attachment of the tail. The 80α connector protein, gp49, forms a tight interaction where each connector monomer binds to two PP monomers in a three-stranded β-sheet (Fig. 4E). Differences in the channel loop between the solution and in situ structures of the 80α PP (Fig. 4D) could likewise reflect a closure of the channel.

During infection, the portal again needs to open to allow for ejection of the DNA. Changes in the baseplate upon interaction with the host are transmitted to the tape measure protein, which is ejected along with the DNA [16]. Apparently, this does not lead to any changes in the portals themselves, since we did not observe any significant differences between the PP in the full and empty capsids (Fig. 4C).

In summary, the 80α portal plays a central role in the phage life cycle, in assembly, DNA packaging, and genome ejection. The interplay between 80α-encoded structural proteins— including PP—and SaPI-encoded factors such as CpmA, CpmB, and TerS is critical to the mechanism of gene transfer and evolution of virulence in this important pathogen. The structures presented here provide further insights into the role of the portal in many of these processes and a basis for interpreting further functional studies.

## ACKNOWLEDGEMENTS

Electron microscopy was done at the UAB Cryo-EM facility (CEMF), supported by the UAB Institutional Research Core Program (IRCP), a UAB Health Science Foundation General Endowment Fund (HSF-GEF) grant to T.D., and National Institutes of Health (NIH) grant S10 OD024978 to T.D. We are also thankful to Dr. Greg Thompson at The University of Alabama for access to their EM facility, while CEMF was unoperational. We acknowledge the assistance of Drs. Thomas Klose and Xueyong Xu at the Midwestern Center for Cryo-Electron Microscopy (MCCEM) at Purdue University with collection of cryo-EM data. MCEEM was supported by NIH grant U24 GM116789 to Dr. Wen Jiang at Purdue University. This work was supported by NIH grant R01 AI083255 to T.D.

## MATERIALS AND METHODS

### 1. Expression and purification of portals

The 80α portal protein gene (ORF42) was amplified by PCR from the 80α genome and cloned into pET21a with a C-terminal hexahistidine tag using the In-Fusion® HD Cloning Kit (Takarabio), yielding pLKP76. For protein expression, the plasmids were transformed into *E.coli* BL21(DE3) cells, grown in LB medium at 37 °C containing 100µg/ml ampicillin, and induced with 1 mM IPTG once the cell density reached A_600_=0.6OD. After induction, cells were grown for 3 hours at 37 °C. The culture was then centrifuged at 10,000 g, and the pellet was stored at −80°C. Cell pellets were resuspended in 15 ml 50 mM HEPES pH 7.4 containing 0.5 M NaCl, 5% glycerol and 20 mM imidazole (binding buffer) in the presence of protease inhibitor cocktail (Sigma-Aldrich). Cells were lysed using an Emulsiflex B15 high pressure disruptor (Avestin) and the lysate was incubated with 1 µl Benzonase® nuclease (EMD Millipore) in the presence of 1 mM MgCl_2_ at 4 °C for 1 hour. The lysate was then centrifuged at 27,000 g for 30 minutes at 4 °C and the supernatant was filtered through a 0.22 µM syringe filter. The filtrate was passed through 1.5 ml of NTA Ni-Agarose (Invitrogen) packed in Econo-Pac® Chromatography Columns (Bio-Rad) equilibrated with binding buffer. Unbound proteins were washed with 20 ml binding buffer and eluted with the same buffer containing 250 mM Imidazole. Purified protein was then dialyzed in 25 mM HEPES buffer pH 7.4, 1M NaCl (dialysis buffer) overnight at 4 °C. The protein was then incubated at 37 °C for 3 to 4 hours to promote oligomerization. Precipitates were removed by centrifugation and the supernatant was concentrated with an Amicon-ultra 50kDa cutoff centrifugal concentrator (Millipore). Concentrated protein was divided into 500 µl aliquots and stored at −80 °C.

### 2. Production of SaPI1 virions

SaPI1 virions were produced by induction of *S. aureus* strain ST65, which contains SaPI1 *tst::tetM* as well as an 80α prophage with a deletion of ORF44 (80ατι*44*) [23]. ORF44 encodes a protein (gp44) that is involved in the survival of phage DNA post injection, but has no effect on SaPI1 replication or assembly [37]. ST65 was grown in CY medium containing 4% β-glycerolphosphate at 32 °C until A_600_ reached 0.5 OD, followed by addition of 0.5 mg mitomycin C per L culture. Growth continued until lysis occurred (5-6 hrs), upon which the lysate was pelleted at 15,000 g for 30 min followed by filtration of the lysate through a 0.22 µm filter. The filtrate was precipitated with 10% PEG 6,000 with 0.5M NaCl at 4°C overnight. The pellet was collected by centrifugation at 15,000 g, 30 min, resuspended in phage buffer (50 mM Tris-HCl pH 7.8, 100 mM NaCl, 4 mM CaCl_2_, 1mM MgSO_4_), and washed with chloroform. CsCl was added to a final concentration of 0.4 g/ml and the solution centrifuged at 389,500 g for 20 hrs. The band containing packaged SaPI1 particles was collected and further purified on a 10-40% sucrose gradient in phage buffer, 154,000 g for 2 hrs. Fractions enriched with SaPI1 particles were collected and pelleted by centrifugation at 165,000 g for 2.5 hours at 4°C. The pellet was resuspended in phage dialysis buffer (20 mM Tris-HCl pH 7.8, 50 mM NaCl, 4 mM CaCl_2_, 1mM MgSO_4_).

### 3. Electron microscopy

Negatively stained samples were made by standard methods: 3 µl of sample was placed on a glow-discharged grid with ultrathin carbon layered on lacey carbon (Electron Microscopy Sciences), washed with three drops of water and stained with 1% uranyl acetate. The grids were observed in an FEI Tecnai F20 microscope operated at 200 kV, equipped with either a Gatan K3 direct electron detector or a Gatan OneView CMOS detector.

For cryo-EM analysis, the PP oligomers were separated at 154,000 g for 2 hrs on a 5-20% sucrose gradient in dialysis buffer. Fractions containing the highest concentration of portal protein oligomers were then dialyzed overnight in 25mM HEPES, pH-7.4, 0.1M NaCl and concentrated using an Amicon Ultra 100 kDa cut-off membrane (Millipore). SaPI1 particles were treated with 1µl Benzonase® nuclease at room temperature for 1 hour prior to cryo-EM grid preparation, followed by dialysis on a 0.025 µm MCE Membrane filter (MF-Millipore) floating on phage dialysis buffer. The buffer was changed three times before collecting the sample.

Cryo-EM samples were prepared using a Vitrobot Mark IV with glow-discharged nickel Quantifoil R2/1 grids or copper C-flat grids, and imaged using an FEI Titan Krios microscope operated at 300 kV and equipped with a Gatan K3 detector and Quantum GIF energy filter at the Midwestern Center for Cryo-Electron Microscopy (MCCEM) at Purdue University. A total of 3,168 images were collected for portals in solution, at a magnification of 81,000 x, corresponding to a pixel size of 1.08 Å, and a total electron dose of 50.97 e^−^/Å^2^. For the SaPI1 virions, 2,796 images were collected at a magnification of 64,000 x (pixel size 1.33 Å), and an electron dose of 35.26 e^−^/Å^2^. See Table S1 for further details.

### 4. Reconstruction of portals

Reconstruction of the *E. coli* expressed portals was performed using RELION-3 [30]. Motion correction was done using RELION’s built-in implementation, and CTF estimation was carried out using GCTF [59]. Images were binned to 4.32 Å/pix; 2D class averages were generated from 2,219 particles picked with Laplacian-of-Gaussian autopicking and used for template-based autopicking of 1,811,063 particles. After rounds of 2D classification, followed by initial model generation with C13 symmetry and 3D classification without symmetry, 96,308 particles were isolated, unbinned and auto-refined with C13 symmetry to 2.8Å resolution. CTF refinement and Bayesian polishing improved the resolution further to 2.4 Å. Local resolution was calculated in RELION-4 with a B-factor of –78.8 Å^2^. (Table S1; Figure S1).

### 5. Reconstruction of SaPI1 capsids

SaPI1 capsids were reconstructed using RELION-4 [30], with motion correction and CTF estimation as above. First, 672 manually picked SaPI1 capsid particles were 2D classified to generate autopicking templates. After autopicking and rounds of 2D classification, particles were classified into 55,513 “full” and 13,774 “empty” capsids, based on the presence or absence of DNA. The two particle groups were processed separately to generate “full” and “empty” capsid reconstructions.

Initial models for full and empty capsids were generated separately without the application of symmetry (C1), followed by 3D classification and refinement. Masked post-processing resulted in maps with a resolution of 4.7 Å and 5.4 Å (FSC=0.143) for the full and empty particles, respectively. Subsequently, an initial model was generated for the full capsids with the application of icosahedral (I1) symmetry, followed by 3D classification and refinement. After post-processing, CTF refinement and Bayesian polishing the final icosahedral reconstruction had a resolution of 3.1 Å (FSC=0.143; Table S1). The empty capsids were processed similarly, resulting in an icosahedrally symmetrical map with a resolution of 3.2 Å (FSC=0.143; Table S1).

### 6. Focused reconstruction of the in situ portal

Focused reconstruction of the in situ portals from full and empty SaPI1 virions was done in RELION-4. Firstly, a reconstruction of the empty virions was generated from 13,500 particles by applying C5 symmetry. This map was then segmented in UCSF ChimeraX [60], and a mask was created to isolate the portal attached to the head-to-tail connector and a part of the tail. The density outside the masked region (corresponding to the capsid) was subtracted from the particles, and an initial model was generated by applying C5 symmetry. Next, the particles underwent 3D classification with C6 symmetry, using the newly generated initial model as a reference. From this step, a subset of 8,414 particles was selected and refined using C12 and C13 symmetries. Individual subunits could be clearly distinguished only in the C12 map, which was therefore chosen for further processing.

This map was subsequently segmented again using a mask that surrounded only the portal. A second signal subtraction was performed with the new mask, followed by initial model generation applying C12 symmetry. After a 3D classification step, a total of 7,423 particles were selected and refined using C12 symmetry, yielding a map with 3.5 Å resolution after post-processing and CTF refinement. Reconstruction of the full virions proceeded similarly, yielding a final reconstruction at 3.2Å resolution using 13,411 particles.

### 7. Reconstruction of the portal-capsid interface

Reconstruction of the portal-capsid interface was done in cryoSPARC [45]. A set of 82,618 full capsid particles were isolated after repicking, and an initial model was generated with icosahedral symmetry. This data set was symmetry expanded according to icosahedral symmetry, followed by 3D classification to identify a subset of particles with tails aligned at a single vertex. The particles were re-oriented to align the tails with the z-axis, re-centered to the portal, and purged of duplicates. After re-extraction, the particles were symmetry expanded according to C5 symmetry and subjected to 3D classification to resolve the symmetry mismatch between the PP and CP. The final subset of 59,457 particles yielded a final reconstruction at 3.4 Å resolution via non-uniform refinement (Table S1; Figure S1).

### 8. Atomic modeling and validation

Atomic model building and refinement was done with Coot [32] and ISOLDE [33] from within ChimeraX [60]. Models were validated in Phenix [61] and MolProbity [62]. The final maps and models were deposited to the EMDB and PDB databases with accession codes EMD-XXXX and PDB IDs YYYY. *PDB and EMDB depositions are in progress and will be finalized before publication.*

## SUPPLEMENTARY INFORMATION

**Supplementary Figure S1.**
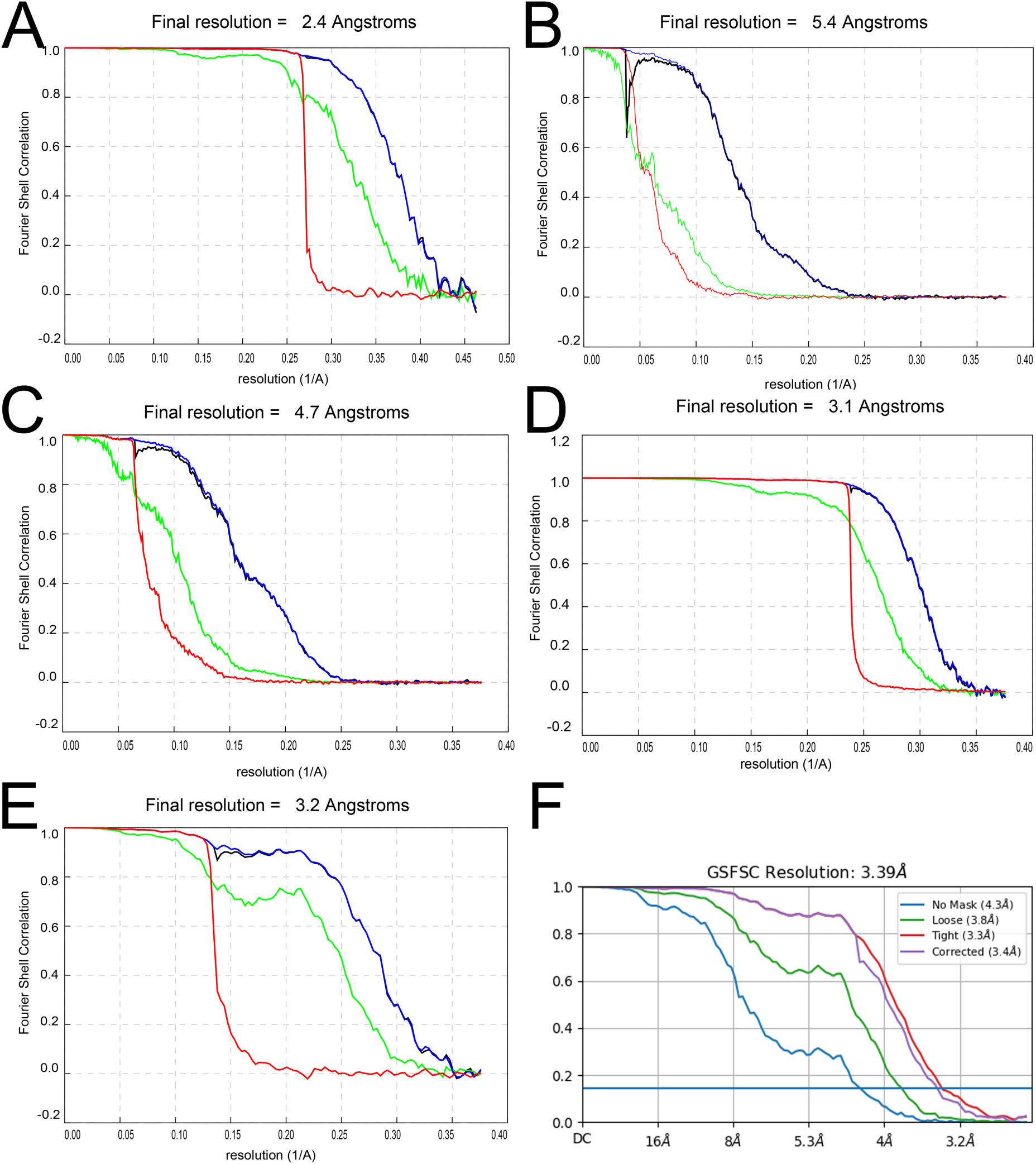
Gold standard Fourier Shell Correlation (FSC) as a function of resolution between two half-maps for each of the reconstructions. **(A)** Portal in solution, C13. **(B)** SaPI1 empty capsid, C1. **(C)** SaPI1 full capsid, C1. **(D)** SaPI1 full capsid, icosahedral. **(E)** In situ portal focused, full capsid, C12. Panels A-E were generated in RELION; for each, the green curve is for the unmasked map, blue is for the masked map, red is for the phase randomized masked maps, and black is the masked curve corrected for the contribution of the mask.. **(F)** FSC for the in situ portal–capsid focused reconstruction from the full capsid, C1, generated in cryoSPARC. The blue, green, purple and red curves are for no mask, loose mask, tight mask, and corrected FSC, respectively.

**Supplementary Figure S2.**
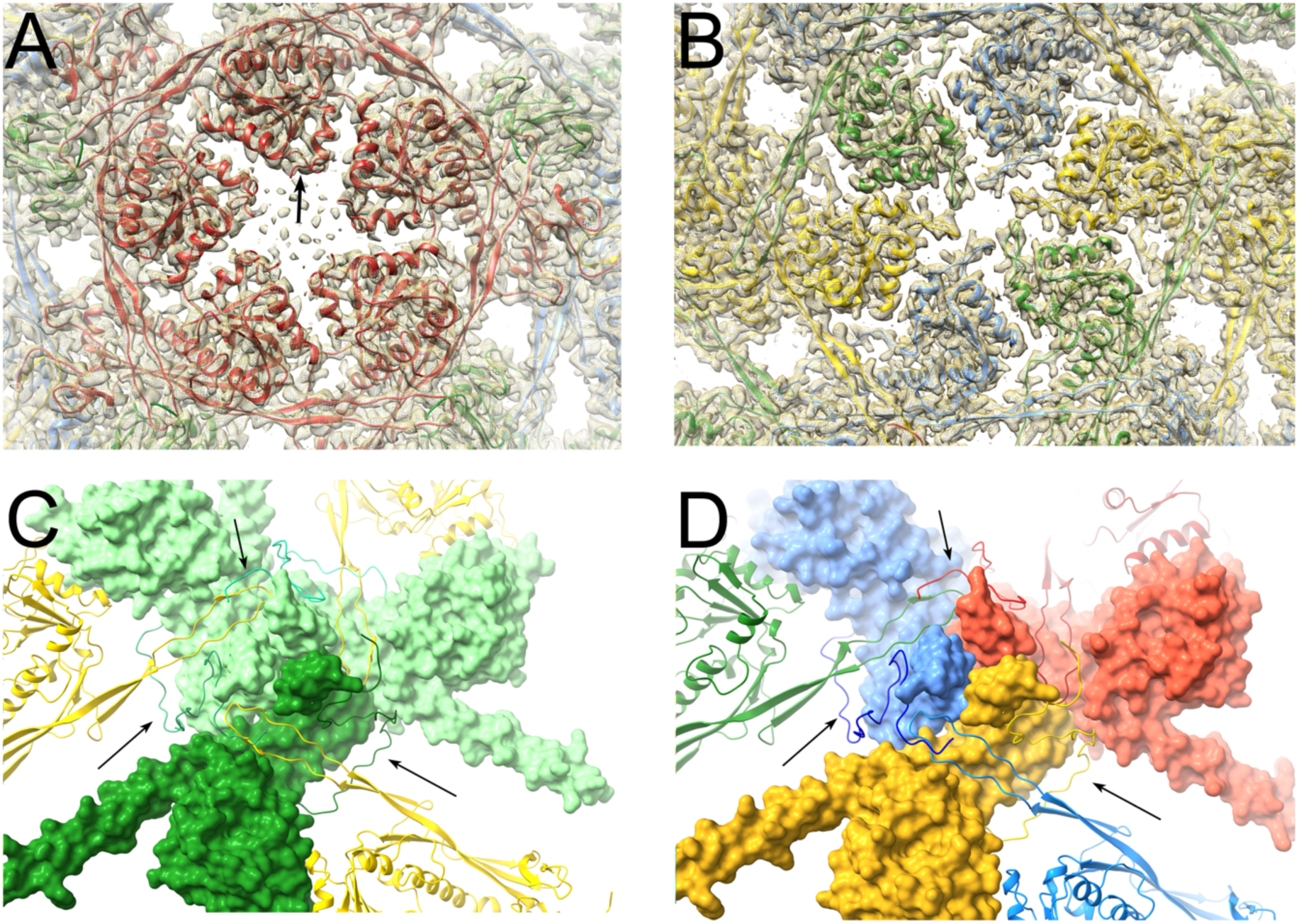
CP interactions. **(A)** View of atomic model (ribbon representation), superimposed on the icosahedral reconstruction density (mesh), viewed down the icosahedral fivefold axis, showing the interaction of A subunit (red) A-domains in the pentamer. One of the missing loops is indicated (arrow). **(B)** View down the icosahedral twofold axis, showing the interaction of A-domains from subunits B (blue), C (green) and D (yellow) in the hexamer. **(C)** Interaction of CP subunits around the icosahedral threefold axis, showing the turret formed by the P loops from three C subunits, shown as a molecular surface. One C subunit is colored a darker shade of green for clarity. The N-arms are shown in ribbon representation and indicated by the arrows. Adjacent D subunits (yellow) are shown in ribbon representation. **(D)** Interaction of CP subunits around the quasi-threefold axis formed by A (red), B (blue) and D (yellow) subunits, shown as a molecular surface. N-arms (arrows) and adjacent subunits are shown in ribbon representation.

**Supplementary Table S1.** Data collection, processing and refinement statistics.

| Structure | Portals in solution | SaPI1 full capsid, C1 | SaPI1 empty capsid, C1 | SaPI1 full capsid, I1 | SaPI1 empty capsid, I1 | PP *in situ* (full) C12 | PP *in situ* (empty) C12 | PP *in situ* (full) C1 |
| --- | --- | --- | --- | --- | --- | --- | --- | --- |
| **Data Collection & Processing** |  |  |  |  |  |  |  |  |
| Microscope |  |  |  | FEI Titan Krios G1 |  |  |  |  |
| Camera |  |  |  | Gatan K3 (counting) |  |  |  |  |
| Energy filter |  |  |  | Gatan Quantum GIF |  |  |  |  |
| Image collection software |  |  |  | Leginon |  |  |  |  |
| Slit width (eV) |  |  |  | 20 |  |  |  |  |
| Voltage (kV) |  |  |  | 300 |  |  |  |  |
| Micrographs collected | 3,168 |  |  |  | 3,581 |  |  |  |
| Defocus range (μm) | 0.7–2.0 |  |  |  | 0.7–2.0 |  |  |  |
| Total exposure (e/Å<sup>2</sup>) | 50.97 |  |  |  | 35.26 |  |  |  |
| Frames per movie | 46 |  |  |  | 38 |  |  |  |
| Pixel size (Å/pix) | 1.08 |  |  |  | 1.33 |  |  |  |
| Reconstruction software | RELION-3.1 | RELION-4.0 | RELION-4.0 | RELION-4.0 | RELION-4.0 | RELION-4.0 | RELION-4.0 | RELION-4.0; CryoSPARC v4.2 |
| Particles extracted | 1,811,063 |  | 97,939 |  |  | 222,708 | 13,500 | 259,743 |
| Final particles | 96,308 | 55,513 | 13,734 | 55,513 | 13,734 | 13,411 | 7,423 | 59,457 |
| Symmetry imposed | C12 | C1 | C1 | I1 | I1 | C12 | C12 | C1 |
| Map resolution (Å, FSC<sub>0.143</sub>) | 2.4 | 3.9 | 4.2 | 3.1 | 3.2 | 3.2 | 3.5 | 3.4 |
| Accession (EMDB) |  |  |  |  |  |  |  |  |
| **Atomic Model Building** |  |  |  |  |  |  |  |  |
| Local refinement software |  |  |  | ISOLDE, Coot |  |  |  |  |
| Global refinement software |  |  |  | Phenix.real_space_refine |  |  |  |  |
| Validation software |  |  |  | Phenix.validation_cryoem |  |  |  |  |
| Sharpening B factor (Å<sup>2</sup>) | -78.80 |  |  | -92.80 | -70.30 | -71.60 | -72.70 | -101.50 |
| Refinement resolution (Å) | 2.5 |  |  | 3.1 | 3.2 | 3.2 | 3.5 | 3.4 |
| Model composition: |  |  |  |  |  |  |  |  |
| Chains | 1 |  |  | 4 | 4 | 2 | 2 | 27 |
| Atoms | 3,900 |  |  | 9,459 | 9,463 | 3,911 | 3,923 | 70,119 |
| Hydrogens | 0 |  |  | 0 | 0 | 0 | 0 | 0 |
| Residues | 476 |  |  | 1,186 | 1,186 | 486 | 478 | 8,633 |
| Waters | 0 |  |  | 0 | 0 | 0 | 0 | 0 |
| Ligands | 0 |  |  | 0 | 0 | 0 | 0 | 0 |
| Bonds (RMSD): |  |  |  |  |  |  |  |  |
| Length (Å) (# > 4σ) | 0.005 (0) |  |  | 0.006 (0) | 0.006 (0) | 0.008 (0) | 0.007 (0) | 0.007 (4) |
| Angles (°) (# > 4σ) | 1.163 (0) |  |  | 1.252 (0) | 1.207 (0) | 1.126 (0) | 1.103 (0) | 1.304 (1) |
| MolProbity score | 0.89 |  |  | 0.91 | 0.91 | 0.64 | 0.83 | 1.09 |
| Clash score | 0.78 |  |  | 0.42 | 0.42 | 0.38 | 1.29 | 1.47 |
| Ramachandran plot (%): |  |  |  |  |  |  |  |  |
| Favored | 97.26 |  |  | 96.17 | 95.83 | 98.13 | 97.89 | 96.77 |
| Allowed | 2.74 |  |  | 3.83 | 4.17 | 1.87 | 2.11 | 3.13 |
| Outliers | 0 |  |  | 0 | 0 | 0 | 0 | 0.11 |
| Rama-Z (Ramachandran plot Z-score, RMSD): |  |  |  |  |  |  |  |  |
| whole (N = 732) | -0.78 (0.34) |  |  | -0.97 (0.23) | -1.14 (0.23) | -0.87 (0.33) | -1.87 (0.32) | -1.44 (0.08) |
| helix (N = 75) | -1.18 (0.29) |  |  | -2.72 (0.23) | -2.82 (0.24) | -0.75 (0.31) | -1.40 (0.29) | -1.46 (0.08) |
| sheet (N = 303) | 1.23 (0.52) |  |  | 1.12 (0.29) | 0.81 (0.29) | 0.58 (0.49) | -0.02 (0.56) | 0.19 (0.12) |
| loop (N = 354) | -0.28 (0.42) |  |  | -0.41 (0.23) | -0.44 (0.25) | -0.75 (0.40) | -1.46 (0.34) | -0.87 (0.09) |
| Rotamer outliers (%) | 0.23 |  |  | 0 | 0 | 0 | 0 | 0.28 |
| C<sub>β</sub> outliers (%) | 0 |  |  | 0 | 0 | 0 | 0 | 0 |
| Peptide plane (%): |  |  |  |  |  |  |  |  |
| Cis proline/general | 0.0/0.0 |  |  | 7.8/0.0 | 7.8/0.0 | 0.0/0.0 | 0.0/0.0 | 4.0/0.0 |
| Twisted proline/general | 0.0/0.0 |  |  | 0.0/0.0 | 0.0/0.0 | 0.0/0.0 | 0.0/0.0 | 0.0/0.0 |
| CaBLAM outliers (%) | 0.21 |  |  | 0.69 | 0.86 | 0.63 | 1.06 | 1.15 |
| Model fit vs. full map: |  |  |  |  |  |  |  |  |
| FSC<sub>0.5</sub> (FSC<sub>0.143</sub>) | 2.5 (2.3) |  |  | 3.2 (2.9) | 3.3 (3.1) | 3.4 (2.8) | 3.7 (3.1) | 3.8 (3.4) |
| CC<sub>box</sub> | 0.34 |  |  | 0.52 | 0.49 | 0.28 | 0.34 | 0.59 |
| CC<sub>mask</sub> | 0.84 |  |  | 0.86 | 0.86 | 0.82 | 0.84 | 0.79 |
| Accession (PDB) |  |  |  |  |  |  |  |  |

**Supplementary Table S2.** Root-mean-square deviation (RMSD) between Cα atoms of portal proteins from different bacteriophages.

| Structure (PDB ID) | 80α PP solution | | 80α PP in situ, full | |
| --- | --- | --- | --- | --- |
|  | RMSD (Å) | Residues§ | RMSD | Residues |
| 80α in situ, full | 0.59 | 320 | – | – |
|  | 7.30 | 461 | – | – |
| 80α in situ, empty | 0.57 | 319 | 0.45 | 453 |
|  | 7.34 | 461 | 0.68 | 466 |
| SPP1 solution (2JES) | 1.25 | 170 | 1.10 | 178 |
|  | 8.18 | 354 | 8.57 | 358 |
| SPP1 in situ (7Z4W) | 1.18 | 195 | 1.16 | 224 |
|  | 9.40 | 417 | 7.24 | 422 |
| P22 in situ, virion (3LJ5) | 1.34 | 24 | 1.36 | 24 |
|  | 17.89 | 392 | 16.08 | 389 |
| Andhra in situ, virion (8EGR) | 1.33 | 18 | 1.04 | 35 |
|  | 20.71 | 289 | 26.04 | 268 |

§The RMSD values are calculated from the number of Cα atom pairs indicated. The first line corresponds to the pruned atom pairs (UCSF Chimera MatchMaker algorithm); the second line corresponds to all atom pairs in the sequences.

